# Revealing the Arabidopsis *AtGRP7* mRNA binding proteome by specific enhanced RNA interactome capture

**DOI:** 10.1101/2024.04.04.588066

**Authors:** Marlene Reichel, Olga Schmidt, Mandy Rettel, Frank Stein, Tino Köster, Falk Butter, Dorothee Staiger

## Abstract

**Background:** The interaction of proteins with RNA in the cell is crucial to orchestrate all steps of RNA processing. RNA interactome capture (RIC) techniques have been implemented to catalogue RNA-binding proteins in the cell. In RIC, RNA-protein complexes are stabilized by UV crosslinking *in vivo*. Polyadenylated RNAs and associated proteins are pulled down from cell lysates using oligo(dT) beads and the RNA-binding proteome is identified by quantitative mass spectrometry. However, insights into the RNA-binding proteome of a single RNA that would yield mechanistic information on how RNA expression patterns are orchestrated, are scarce.

**Results:** Here, we explored RIC in Arabidopsis to identify proteins interacting with a single mRNA, using the circadian clock-regulated *Arabidopsis thaliana* GLYCINE-RICH RNA-BINDING PROTEIN 7 (*AtGRP7*) transcript, one of the most abundant transcripts in Arabidopsis, as a showcase. Seedlings were treated with UV light to covalently crosslink RNA and proteins. The *AtGRP7* transcript was captured from cell lysates with antisense oligonucleotides directed against the 5’untranslated region (UTR). The efficiency of RNA capture was greatly enhanced by using locked nucleic acid (LNA)/DNA oligonucleotides, as done in the enhanced RIC protocol. Furthermore, performing a tandem capture with two rounds of pulldown with the 5’UTR oligonucleotide increased the yield. In total, we identified 356 proteins enriched relative to a pulldown from *atgrp7* mutant plants. These were benchmarked against proteins pulled down from nuclear lysates by *AtGRP7 in vitro* transcripts immobilized on beads. Among the proteins validated by *in vitro* interaction we found the family of Acetylation Lowers Binding Affinity (ALBA) proteins. Interaction of ALBA4 with the *AtGRP7* RNA was independently validated via individual-nucleotide resolution crosslinking and immunoprecipitation (iCLIP). The expression of the *AtGRP7* transcript in an *alba* loss-of-function mutant was slightly changed compared to wild-type, demonstrating the functional relevance of the interaction.

**Conclusion:** We adapted specific RNA interactome capture with LNA/DNA oligonucleotides for use in plants using *AtGRP7* as a showcase. We anticipate that with further optimization and up-scaling the protocol should be applicable for less abundant transcripts.

## Background

The interaction of proteins with RNA in the cell is crucial to orchestrate all steps of RNA processing including splicing, nuclear export and decay [1–3]. The identification of RNA-binding proteins (RBPs) interacting with an RNA thus provides insights into RNA biogenesis and function. Research towards assembling a global inventory of proteins binding to RNA *in vivo* has been enabled by recent technological advancements increasing the sensitivity in mass spectrometry (MS). In RNA Interactome Capture (RIC) approaches, RNA and associated proteins are covalently linked by irradiation with 254 nm UV light. This creates radicals of the nucleobases that react with amino acids in their immediate vicinity. The polyadenylated RNA fraction is then recovered by hybridization with bead-coupled oligo(dT) under stringent conditions. Associated proteins are eluted by ribonuclease digestion and subjected to liquid chromatography/tandem MS, leading to the identification of about 800 mRNA-bound proteins in HeLa and HEK293 cells [4, 5]. Subsequently, RIC was applied to different mammalian cell lines and tissues, and to a range of organisms including yeast [6–8], *Caenorhabditis elegans* [6] or *Drosophila melanogaster* [9, 10], reviewed in [8, 11].

The sensitivity of RIC was increased through the use of locked nucleic acids (LNAs) in the “enhanced RIC” (eRIC) protocol. In LNA nucleosides, a methylene bridge “locks” the ribose between 2’-O and 4’-C, favoring Watson-Crick base-pairing and increasing the stability of duplexes and resistance to nucleases [12]. Furthermore, the oligonucleotides are covalently linked to magnetic beads, thus tolerating more stringent salt conditions and the use of chaotropic detergents employed to reduce non-specific interactions and increase signal-to-noise ratios.

The RIC protocol was also applied to *Arabidopsis thaliana*, requiring major adjustments to the challenges posed by plant tissue [13–15]. As plants contain UV absorbing pigments, the UV dosage for RNA-protein crosslinking in the tissue was increased. Moreover, stringent conditions for lysis were applied to overcome the hurdles by the rigid cell wall and the inherently higher RNase content of plant tissue (for review, see [16–18]). The RNA-binding proteome was profiled in different Arabidopsis tissues including leaves of four-week-old plants [13], leaf mesophyll protoplasts [14], etiolated seedlings [15], cell cultures derived from roots [13] or an egg-cell like callus [19]. Moreover, Bach-Pages improved the purification conditions to more efficiently remove contaminations during oligo(dT) pulldown [20].

Collectively, these RIC approaches greatly advanced our understanding about the types of proteins capable of binding RNA, including many proteins lacking conventional RNA-binding domains or metabolic enzymes (reviewed in [11]). However, insights into the RBPome of a single RNA and its dynamic changes, which would yield mechanistic information on how RNA expression patterns are orchestrated, are scarce. Until recently, the identification of proteins interacting with a defined RNA or candidate regulatory regions within an RNA were mainly identified *in vitro* by immobilizing the RNA bait, incubating it with protein extracts from the tissue of choice and retrieving bound proteins for MS. For this, RNA can be chemically synthesized or *in vitro* transcribed in the presence of biotinylated UTP, allowing loading onto streptavidin beads [21]. Alternatively, RNAs can be spiked with a recognition sequence for a high affinity RBP such as the coat protein of phage MS2 [22]. Upon incubation of the MS2-tagged RNA with protein extracts of the tissue of choice, the assembled complexes are captured with immobilized MS2 coat protein-GFP and proteins co-purifying with the RNA are identified via MS [23]. To retrieve proteins regulating processing of miRNA precursors, a 16-nucleotide hairpin that is bound with high affinity by the cleavage deficient Csy4* Cas nuclease has been used as tag. The Csy4* protein is first immobilized onto beads followed by loading of an *in vitro* transcript comprising the miRNA precursor tagged with the hairpin sequence. After incubation with nucleoplasmic or cellular extracts, the protein-RNA complexes are retrieved by pulling down the Csy4*. Proteins are released by reactivation of the Csy4* cleavage activity and then subjected to MS [24, 25].

Overall, these *in vitro* approaches do not take into account features of the native environment including epitranscriptomic modifications of binding sites or interactions of RBPs with other molecules in the cell which may affect binding. Furthermore, the *in vitro* structure may not recapitulate the secondary structure found in the cell. Therefore, methods to capture proteins interacting with a single RNA species *in vivo* are needed.

Zooming in on a single RNA *in vivo* has been achieved for the 3.7 kb long human long noncoding RNA (lncRNA) NEAT1 (nuclear-enriched abundant transcript 1) and the 8 kb lncRNA MALAT1 (metastasis-associated lung adenocarcinoma transcript 1) [26, 27]. West and co-workers extended the Capture Hybridization Analysis of RNA Targets (CHART) procedure, originally designed to isolate lncRNAs bound to DNA from formaldehyde crosslinked tissue to identify sites of lncRNA interaction with genomic loci, to proteomics analysis by MS (CHART-MS) [28, 29]. Subsequent studies focused on the development of methods to comprehensively profile proteins interacting with the X-inactive specific transcript (Xist), a long noncoding RNA 17 kb in length required for X-chromosome inactivation and thus dosage compensation in female placental mammals. In a comprehensive identification of RBPs by mass spectrometry (ChIRP-MS), Chu and coworkers employed long biotinylated antisense oligonucleotides (AOs) spanning the entire Xist transcript. After formaldehyde fixation of the tissue and preparation of lysates, these tiling AOs were hybridized to Xist under stringent conditions. Oligonucleotide-bound RNA and the *in vivo* assembled proteins were efficiently pulled down with streptavidin magnetic beads [30]. ChIRP-MS was also employed to recover proteins associated with human U1 and U2 snRNAs and Arabidopsis U1 snRNA [30, 31]. In RNA antisense purification (RAP)-MS, McHugh and co-workers used UV light for crosslinking, followed by hybridization with long biotinylated AOs under strong denaturing conditions [32]. Mass spectrometry identified a suite of RBPs that were functionally linked to chromatin spreading and silencing of Xist transcription [30, 32]. Pulldown with AOs designed against the lncRNA Pluripotency and Hepatocyte Associated RNA Overexpressed in HCC (PHAROH) identified the translational repressor TIAR that binds to a 71 nucleotide hairpin in PHAROH, leading to increased translation of MYC [33].

In Arabidopsis, Crespi and colleagues searched for proteins interacting with the *ALTERNATIVE SPLICING COMPETITOR* (*ASCO*) lncRNA, which has been shown to regulate the activity of the splicing regulator NUCLEAR SPECKLE RNA-BINDING PROTEIN a (NSRa) [34]. Pulldown with a set of 20-nucleotide long biotinylated AOs covering the length of the lncRNA identified the splicing factor PRP8, in line with the model of *ASCO* as a competitive inhibitor of the action of splicing factors [35].

A major drawback of these tiling approaches lies in the fact that they cannot distinguish between RNA isoforms or between closely related members of multigene families that are particularly prevalent in Arabidopsis. To increase the specificity, Rogell and colleagues modified the enhanced RIC protocol, using specific LNA/DNA AOs targeted at defined regions of a transcript and scrambled probes as control [36]. This “specific ribonucleoprotein (RNP) capture” procedure was tested against HeLa cell 18S and 28S ribosomal RNA and uncovered proteins previously unknown to interact with rRNAs or to be involved in rRNA biology.

Here, we explored RNA interactome capture in Arabidopsis to identify proteins interacting with a single mRNA that may contribute to processing and function of this mRNA. We chose the circadian-clock regulated transcript *AtGRP7* encoding a glycine-rich RNA binding protein as a paradigm, as it is one of the most abundant transcripts in Arabidopsis [37, 38]. In addition to transcriptional regulation by the circadian clock, *AtGRP7* is regulated at the posttranscriptional level. Elevated *At*GRP7 levels entail alternative splicing at a cryptic splice site in the intron leading to retention of the first half of the intron including a premature termination codon and, consequently, degradation via the Nonsense mediated decay pathway [39–42]. *At*GRP7 binds to its own pre-mRNA *in vitro* and *in vivo* [42–44] and mathematic modelling unveiled that this posttranscriptional regulation indeed contributes to the *AtGRP7* oscillations [45]. We hypothesize that additional proteins binding to the *AtGRP7* transcript *in vivo* contribute to shaping its temporal expression pattern.

An antisense oligonucleotide directed against the 5’ untranslated region (UTR) proved most efficient in capturing *AtGRP7* transcript from lysates of UV crosslinked plants. Furthermore, the use of LNA oligonucleotides greatly enhanced the capture efficiency. The largest number of proteins were identified by two rounds of pulldown with the specific 5’UTR LNA oligonucleotide. Performing a tandem capture approach with the specific 5’UTR LNA oligonucleotide first followed by capture with an LNA oligo(dT) oligonucleotide led to an increased purity of the captured *AtGRP7* transcript but came at the cost of a reduced number of proteins detected in mass spectrometry. The proteins identified by the *in vivo* interactome captures were benchmarked against proteins recovered by *in vitro* pulldown using biotinylated, bead immobilized *AtGRP7* UTRs. Among the proteins interacting with the *AtGRP7* transcript *in vitro* was the family of ACETYLATION LOWERS BINDING AFFINITY (ALBA) proteins. Interaction of ALBA4 with the *AtGRP7* RNA was independently validated via individual-nucleotide resolution crosslinking and immunoprecipitation (iCLIP).

Overall, we adapted RNA interactome capture with LNA/DNA oligonucleotides to identify the complement of proteins interacting with a single RNA in plants. We anticipate that further optimization and up-scaling of the protocol for transcripts of interest along with the continuous improvement of mass spectrometry sensitivity will further increase the detection proteins that are enriched relative to interactome capture from control plants devoid of the transcript of interest.

## Methods

### Plant lines

The *grp7-1* T-DNA mutant and the *AtGRP7::AtGRP7-GFP* plants expressing an *At*GRP7-GFP fusion under control of 1.4 kb of the *At*GRP7 promoter and the *At*GRP7 5’UTR, intron and 3’UTR in the *grp7-1* T-DNA mutant have been described before and are available from Dorothee Staiger [43, 46–48]. The *alba456* triple mutant, the transgenic line expressing the ALBA4-GFP fusion protein driven by 1.7 kb of the *ALBA4* promoter in the *alba456* mutant background, and the 35S-GFP control line expressing GFP under control of the Cauliflower Mosaic Virus 35S RNA promoter have been described and are available from Anthony Millar, ANU Canberry, Australia [49].

### Plant growth and UV crosslinking

Seeds were sterilized by chlorine gas (100 ml bleach and 3 ml 37% HCl) for 4 hours in a sealed desiccator jar and germinated on half-strength Murashi Skoog plates [50]. Seedlings were grown in 16 h light-8 h dark cycles at 20°C for 14 days. UV crosslinking was performed at zt13 (zeitgeber time 13, 13 h after lights on) at 2000 mJ/cm^2^ [43, 51].

### Probe design

LNA antisense oligonucleotides for *AtGRP7* were designed using the QIAGEN LNA Oligo Optimizer tool. Probes were designed to minimize self-complementarity of the sequence, secondary structure and no complementarity to other Arabidopsis transcripts, in particular the *AtGRP8* paralog. Furthermore, complementary regions in the *AtGRP7* transcript were selected that have little secondary structure as determined by the Vienna RNA fold server. Probes were 18-22 nts long with a melting temperature of 68-84°C and contained a flexible C6 linker carrying a primary amino group at the 3’ end to couple them covalently to magnetic beads. All probes were HPLC purified. LNA2.T are (dT)_20_ oligonucleotides with every second dT substituted by an LNA-thymine [52].

### Coupling of LNA oligonucleotides to magnetic beads

Probes were coupled to carboxylated magnetic beads (M-PVA C11, PerkinElmer, Waltham, MA, USA) in DNA/RNA low binding tubes. The oligonucleotides were resuspended in nuclease-free water (100 µM final concentration). The bead slurry (50 mg/ml) was washed three times with 5 volumes of 50 mM 2-(N-morpholino)ethanesulfonic acid (MES) pH 6.0. One volume of prewashed bead slurry was combined with 5 volumes of freshly prepared 20 mg/ml *N*-(3-dimethylaminopropyl)-*N′*-ethylcarbodiimide hydrochloride (EDC-HCl) in MES buffer and 0.66 volumes of the oligonucleotide solution (e.g. for 150 μl of bead slurry, 750 μl EDC-HCL solution and 10 nmol of probe in 100 μl H_2_O were added). Coupling was performed for 5 h at 50°C in a thermomixer (800 rpm). To monitor the coupling efficiency, the oligonucleotide concentration was measured in the supernatant by NanoDrop. Subsequently, the beads were washed 3 times with 2 ml 1x PBS pH 7.4 and residual free carboxyl residues were inactivated by adding 2 ml of 200 mM ethanolamine pH 8.5 for 1 h at 37°C and 800 rpm in the thermomixer. The coupled beads were finally washed 3 times with 2 ml 1 M NaCl and stored in 0.1% (v/v) PBS-Tween at 4°C until use.

### Specific RNA interactome capture *in vivo* (medium- and large-scale experiments)

Above-ground tissue collected from 14 day old, UV crosslinked *grp7-1* or GRP7-GFP seedlings, respectively, was ground into a fine powder with liquid nitrogen. The following steps are described for 8 g of ground tissue, which were split up into two 50 ml tubes with 4 g each and resuspended in 25 ml lysis buffer (50 mM Tris-HCl pH 7.5, 500 mM NaCl, 0.5% SDS [w/v], 1 mM EDTA) supplemented with 1x protease inhibitor (complete EDTA free, Roche), 2.5% [w/v] polyvinylpyrrolidone 40 (PVP40), 1% beta-mercaptoethanol, 5 mM DTT and 10 mM Ribonucleoside Vanadyl Complex. The lysate was centrifuged for 12 min at 5000 rpm and 4°C. The cell extract was then passed through a 0.45 μm filter and subsequently through a 27G needle to shear genomic DNA. In a fresh 50 ml tube, 42.5 ml cell extract were mixed with 7.5 ml formamide (15% (v/v) final concentration), and an aliquot of 100 μl was saved for RNA and protein analysis. For pre-clearing, 600 ml M-PVA C11 beads were equilibrated by washing three times in 3 volumes of lysis buffer and then added to 50 ml cell extract. After rotation for 1h at 4°C, the tubes were placed on a magnet for about 15 minutes at 4°C and the cell extract was transferred to a new tube. For the specific RNP capture, 600 ml M-PVA C11 beads coupled to 40 nmol *AtGRP7* 5’UTR LNA oligo were washed three times in 3 volumes of lysis buffer, resuspended in 500 µl lysis buffer and heated up to 90°C for 3 min to relax secondary structures. The beads were then added to 50 ml of cell extract and incubated for 2 h at 4°C on a rotator. Tubes were then placed on a magnet and if a second round of capture was performed, the cell extract was transferred to a new tube and a fresh aliquot of beads coupled to the *AtGRP7* 5’UTR LNA oligo was added and incubated for 2 h at 4°C with rotation. After hybridization and magnetic separation, a 100 μl aliquot of the cell extract was taken for RNA and protein analysis and the rest was discarded. Beads were resuspended in 2 ml of lysis buffer and transferred to a 2 ml DNA/RNA low binding tube. Beads were then washed four times in 2 ml 2 x SSC buffer supplemented with 0.5% SDS and 5 mM DTT followed by one wash with 2 ml 1 x SSC buffer (without SDS and DTT). All washed were carried out for 5 min at room temperature on a rotator. For pre-elution of unspecific binders, the beads were resuspended in 600 μl H_2_O and incubated for 5 min at 40°C and 800 rpm. After magnetic separation, the pre-eluates were saved for RNA and protein analysis. RNA-protein complexes were finally heat-eluted by resuspending the beads in 600 μl elution buffer (20 mM Tris-HCl pH 7.5, 1 mM EDTA) and incubation for 3 min at 90°C and 800 rpm. Tubes were placed on a magnet, the eluates were transferred to a new tube and an aliquot was saved for RNA analysis. The rest of the eluate was either used for RNase treatment, protein concentration and western blot or mass spec, respectively, or subjected to another purification by oligo(dT) or LNA2.T capture.

For the medium-scale capture, this procedure was carried out with 32 g of ground tissue (8 x 4g in 50 ml tubes) and for the large-scale capture, 100 g of ground tissue were used (24 x 4g in 50 ml tubes) and the pre-eluates and eluates were pooled.

The preliminary test of capture efficiency of different LNA oligonucleotides was performed in the same way, except that only 12.5 ml of cell extract and 150 μl of beads coupled to 10 nmol of the respective LNA oligonucleotides were used.

### LNA2.T capture

The pooled eluate from the capture with the *AtGRP7* 5’UTR LNA oligonucleotide (14.4 ml) was aliquoted into twelve 5 ml tubes with 1200 μl eluate each. This was mixed with 2667 μl elution buffer, 833 μl 3 M NaCl (final concentration of 0.5 M) and 300 μl of M-PVA C11 beads coupled to 20 nmol of LNA2.T oligo, which were first washed three times in 3 volumes of lysis buffer, resuspended in 300 μl lysis buffer and heated to 90°C for 1 min. Hybridization was carried out for 2 h at 4°C on a rotator. After magnetic separation for 10 min at 4°C, the supernatant was transferred into a new tube and a second round of capture was performed with a fresh aliquot of LNA2.T coupled beads. Washes, pre-elution and elution were performed as described above, except that pre-elution and elution were carried out in a volume of 300 μl. Pre-eluates and eluates were pooled and used for quality controls and mass spec.

### Recycling of beads coupled to LNA oligos

Beads coupled to LNA oligonucleotides can be reused multiple times and were recycled by resuspending them in 1 mL of nuclease-free water and incubation for 5–10 min at 95°C and 800 rpm. Immediately afterwards, before bead slurry cool down, beads were collected by magnetic force, and the supernatant discarded. Beads were then washed 3 times with 5 volumes of water and 3 times with 5 volumes of lysis buffer and stored in 0.1% PBS–Tween at 4°C until use. To avoid cross-contamination, beads were only reused for the same line.

### Oligo(dT) capture

The pooled eluate from the capture with the *AtGRP7* 5’UTR LNA oligo (2400 μl) was aliquoted into four 5 ml tubes with 600 μl eluate each. This was mixed with 3650 μl elution buffer (20 mM Tris-HCl (pH 7.5), 1 mM EDTA) and 500 μl 5 M LiCl (final concentration of 0.5 M). 240 μl of oligo(dT) bead slurry (New England Biolabs, Frankfurt, Germany) were washed three times in 1 mL of lysis buffer (20 mM Tris-HCl pH 7.5, 500 mM LiCl, 1 mM EDTA, 5 mM DTT, 0.5% (w/v) LiDS), resuspended in 250 μl elution buffer and then added to the eluate. Hybridization was carried out for 2h at 4°C on a rotator. After magnetic separation for 5 min at 4°C, the supernatant was removed, and the beads were resuspended in lysis buffer and transferred to a 2 mL tube. Beads were then washed twice with 2 mL of buffer I (20 mM Tris-HCl pH 7.5, 500 mM LiCl, 1 mM EDTA, 5 mM DTT, 0.1% (w/v) LiDS), buffer II (20 mM Tris-HCl pH 7.5, 500 mM LiCl, 1 mM EDTA, 5 mM DTT) and buffer III (20 mM Tris-HCl pH 7.5, 200 mM LiCl, 1 mM EDTA, 5 mM DTT). Each wash was performed for 5 min at 4°C on a rotator. To elute RNA-protein complexes, beads were resuspended in 400 µl elution buffer and incubated for 3 min at 50°C and 800 rpm. Eluates were pooled and used for quality controls and mass spec.

### Quality controls

#### RNA extraction, cDNA synthesis and qPCR

RNA was isolated from the input and the supernatant using commercial TRIzol (Thermo Fischer, Waltham, MA, USA) according to the manufacturer’s instructions and RNA integrity was checked by denaturing agarose-formaldehyde gel electrophoresis. 4 µl of the RNA from input and supernatant and 7 µl of the pre-eluates and eluates were digested with RQ1 DNase I (Promega, Walldorf, Germany) and cDNA synthesis was carried out with Superscript IV reverse transcriptase (Thermo Fischer) and random hexamer primers according to the manufacturer’s instructions. qPCR was performed in a volume of 10 μL with the iTaq SYBR GREEN supermix (Biorad, Hercules, CA, USA) using 45 cycles of 15 s at 95°C and 30 s at 60°C in a CFX96 cycler (Biorad). Primers are listed in Additional file 1.

#### RNase treatment, protein concentration and silver staining

The remainder of the pre-eluate and eluate was treated with RNase cocktail (Thermo Fischer) (0.5 µl per 1200 µl eluate/pre-eluate) for 1 h at 37°C. After concentration using an Amicon filter device with a 3K cut-off, the samples were subjected to SDS-PAGE using a 4-12% NuPAGE Bis-Tris gel run at 100 V in 1x MOPS buffer followed by silver staining as described in Reichel et al. [15].

### Immunoblot analysis

Immunoblot analysis was done essentially as described in Meyer at el. [43]. Primary antibodies were a polyclonal antibody against HISTONE 3 (Agrisera, Vännäs, Sweden, AS10710; rabbit; dilution 1:5000), and a monoclonal antibody against GFP (Roche, catalog number 11 814 460 001; mouse; dilution 1:1000). Secondary antibodies were HRP-coupled anti-rabbit IgG (Sigma-Aldrich catalog number A 0545; dilution 1:5000) or HRP-coupled anti-mouse IgG (Sigma-Aldrich catalog number A0168; dilution 1:2500). Chemiluminescence detection of the immunoblots was done with Fusion-FX6 system (Vilber Lourmat, Eberhardzell, Germany).

### Mass spectrometry

#### Sample preparation

Reduction of disulphide bridges in cysteine containing proteins was performed with dithiothreitol (56°C, 30 min, 10 mM in 50 mM HEPES, pH 8.5). Reduced cysteines were alkylated with 2-chloroacetamide (room temperature, in the dark, 30 min, 20 mM in 50 mM HEPES, pH 8.5). Samples were prepared using the SP3 protocol [53, 54] and trypsin (sequencing grade, Promega) was added in an enzyme to protein ratio 1:50 for overnight digestion at 37°C. Next day, peptide recovery in HEPES buffer by collecting supernatant on magnet and combining with second elution wash of beads with HEPES buffer.

Peptides were labelled with TMT6plex [55] Isobaric Label Reagent (ThermoFisher) according the manufacturer’s instructions. Samples were combined for multiplexing and for further sample clean up an OASIS® HLB µElution Plate (Waters) was used. Offline high pH reverse phase fractionation was carried out on an Agilent 1200 Infinity high-performance liquid chromatography system, equipped with a Gemini C18 column (3 μm, 110 Å, 100 x 1.0 mm, Phenomenex, Aschaffenburg, Germany) [15].

#### LC-MS/MS

An UltiMate 3000 RSLC nano LC system (Dionex) fitted with a trapping cartridge (µ-Precolumn C18 PepMap 100, 5µm, 300 µm i.d. x 5 mm, 100 Å) and an analytical column (nanoEase™ M/Z HSS T3 column 75 µm x 250 mm C18, 1.8 µm, 100 Å, Waters). Trapping was carried out with a constant flow of trapping solution (0.05% (v/v) trifluoroacetic acid in water) at 30 µL/min onto the trapping column for 6 minutes. Subsequently, peptides were eluted via the analytical column running solvent A (0.1% (v/v) formic acid in water, 3% (v/v) DMSO) with a constant flow of 0.3 µL/min, with increasing percentage of solvent B (0.1% (v/v) formic acid in acetonitrile, 3% DMSO (v/v)). The outlet of the analytical column was coupled directly to an Orbitrap Fusion™ Lumos™ Tribrid™ Mass Spectrometer (Thermo Fischer) using the Nanospray Flex™ ion source in positive ion mode. The peptides were introduced into the Fusion Lumos via a Pico-Tip Emitter 360 µm OD x 20 µm ID; 10 µm tip (CoAnn Technologies) and an applied spray voltage of 2.4 kV. The capillary temperature was set at 275°C. Full mass scan (MS1) was acquired with mass range 375-1500 m/z in profile mode in the orbitrap with resolution of 60000. The filling time was set at maximum of 50 ms. Data dependent acquisition (DDA) was performed with the resolution of the Orbitrap set to 15000, with a fill time of 54 ms and a limitation of 1×105 ions. A normalized collision energy of 36 was applied. MS2 data was acquired in profile mode.

#### Data analysis

IsobarQuant [56] and Mascot (v2.2.07) were used to process the acquired data, which was searched against a Uniprot *Arabidopsis thaliana* proteome database containing common contaminants and reversed sequences. The following modifications were included into the search parameters: Carbamidomethyl (C) and TMT10 (K) (fixed modification), Acetyl (Protein N-term), Oxidation (M) and TMT10 (N-term) (variable modifications). For the full scan (MS1) a mass error tolerance of 10 ppm and for MS/MS (MS2) spectra of 0.02 Da was set. Further parameters were set: Trypsin as protease with an allowance of maximum two missed cleavages: a minimum peptide length of seven amino acids; at least two unique peptides were required for a protein identification. The false discovery rate on peptide and protein level was set to 0.01.

The raw output files of IsobarQuant (protein.txt – files) were processed using the R programming language [57]. Contaminants were filtered out and only proteins that were quantified with at least two unique peptides were considered for the analysis. 18 (Fig. 3D), 64 (Fig. 4A), 387 (Fig. 4C) and 2626 (Fig. 6E) proteins passed these criteria. Raw TMT reporter ion intensities (‘signal_sum’ columns) were used for further analysis. For Fig. 3D and 4C, GRP7-GFP and *grp7-1* fold-changes were calculated on raw TMT reporter ion intensities. For Fig. 4A, raw TMT reporter ion intensities were normalized using vsn before (variance stabilization normalization, [58]). For Fig. 6E, raw TMT reporter ion intensities were first cleaned for batch effects using limma [59] and further normalized using vsn [58]. Proteins were tested for differential expression using the limma package. The replicate information was added as a factor in the design matrix given as an argument to the ‘lmFit’ function of limma. A protein was annotated as a hit with a false discovery rate (fdr) smaller 0.05 and a fold-change of at least 2-fold.

### *In vitro* pulldowns

#### Isolation of nucleoplasmic proteins

14-day old Col-0 seedlings were snap-frozen in liquid nitrogen and ground into a fine powder by mortar and pestle. 5 g of ground material were resuspended in 15 mL lysis buffer (20 mM Tris-HCl pH 7.5, 25% (v/v) glycerol, 20 mM KCl, 2 mM EDTA, pH 8.0, 2.5 mM MgCl_2_, 250 mM sucrose, 5 mM DTT) [24]. The suspension was filtered through a 100 µm mesh and 2 layers of miracloth. The sample was diluted to 50 mL with lysis buffer and centrifuged at 1500 g for 30 min at 4 °C. The crude nuclear pellet was resuspended in 12 ml NRBT (20 mM Tris-HCl pH 7.4, 25% (v/v) glycerol, 2.5 mM MgCl_2_, 0.2% Triton X-100), incubated on ice for 3 min and centrifuged at 1500 g for 30 min at 4°C. This washing step was repeated 5 to 6 times until the pellet was whitish. Finally, the pellet was suspended in 3 volumes (~600 μL) of protein extraction buffer (20 mM Hepes-KOH, pH 7.5, 5% (v/v) glycerol, 1.5 mM MgCl_2_, 0.2 mM EDTA pH8.0, 420 mM NaCl, 5 mM DTT, 1x Complete Protease inhibitor (Roche, San Francisco, CA, USA) [60] and adjusted to 150 mM NaCl.

#### *In vitro* transcription

The template for the 5’UTR was PCR amplified from a plasmid harbouring the *AtGRP7* 5’UTR and 15 nucleotides of exon 1 driven by the T7 promoter using primers gB_5’UTR_t7_fwd and gB_5’UTR_rev (Additional file 1). The template for the 3’UTR was PCR amplified from a plasmid harbouring the *AtGRP7* 3’UTR driven by the T7 promoter using primers gB T7 GRP7 3’UTR fwd and gB T7 GRP7 3’UTR rev (Additional file 1). PCR products were purified using the NucleoSpin Gel and PCR Clean-up (Macherey-Nagel) and used for *in vitro* transcription.

*In vitro* transcription was performed using the MEGAshortscript T7 transcription kit (Invitrogen, Carlsbad, CA, USA) according to the manufacturer’s instructions with a mixture of Biotin 11-UTP and UTP so that each transcript contained on average three biotinylated uracil. The *in vitro* transcripts were purified using the RNA Clean & Concentrator-25 Kit (Zymo Research, Freiburg, Germany) and analyzed via Urea-PAGE.

#### Coupling of *in vitro* transcripts to magnetic beads

50 pmol of the biotinylated *in vitro* transcripts were diluted in 2x folding buffer (40 mM Tris-HCl pH 7.5, 200 mM NaCl, 6 mM MgCl_2_), incubated at 70°C for 10 min, put on ice for 1 min and incubated at room temperature for 10 min. An aliquot was taken as input. Thirty μl of magnetic Dynabeads MyOne Streptavidin beads were washed in 1x folding buffer and incubated with the renatured *in vitro* transcripts at 4°C for 1.5 to 2 h. Beads were placed on a magnet and an aliquot of the supernatant was taken to monitor coupling efficiency. Additionally, 5% of the beads were used to elute the RNA in formamide containing loading dye (90% formamide, 10% glycerol, bromophenol blue, xylene cyanol) and check the RNA integrity on the beads before pulldown by urea PAGE.

#### Capture of RNA-protein complexes

The rest of the beads was incubated with 2 mL (ca. 800-1000 μg) nucleoplasmic protein extract for 2 h at 4°C in an end-over-end rotator. Subsequently, beads were placed on a magnet, an aliquot of the supernatant was taken for protein analysis and the rest was discarded. The beads were washed three times with 500 μl protein wash buffer (PWB, 20 mM Tris-HCl pH 7.5, 100 mM NaCl, 5% (v/v) glycerol, 1 mM TCEP, 0.1% Triton X-100) for 5 min at 4°C and 800 rpm. 5% of the beads were used to elute the RNA with formamide containing loading dye and check the RNA integrity after the pulldown by urea PAGE. Proteins were eluted from the remaining 95% of the beads by resuspending them in 40 μl 1x LDS sample buffer (Thermo) and incubation for 10 min at 70°C. Proteins were either analysed by silver staining or mass spectrometry. A pulldown with empty beads was performed side-by-side as a negative control. Four replicates were processed in parallel and analysed by mass spec.

#### Mass spectrometry

For the pulldown with the 5’UTR, samples were processed as described for the *in vivo* pulldowns. The 3’UTR samples were measured with an EasyLC/Q Exactive Plus mass spectrometer platform as described [24]. The data was analysed as described for the *in vivo* pulldown. Logarithmic transformations of the enrichment were plotted against the determined *p*-value. Proteins with a *p*-value ≤ 0.05 and a log_2_ fold change ≥ 1.0 were considered as significantly enriched.

### GO term analysis

Genes enriched in gene ontology (GO) terms were analyzed in the online Thalemine database using the default background population and Holm–Bonferroni test correction.

### Circadian time course

Col-0 wild-type plants and the *alba456* mutant were grown on half strength MS medium in entraining long day conditions at 20°C for 12 days before shifting them to continuous light (LL). Plants were harvested at two-hour intervals in long days and from 28 to 76 h after transfer to LL. Plants were quick frozen in liquid nitrogen and RNA was isolated using the Universal RNA purification kit (Roboklon, Berlin, Germany). cDNA synthesis was performed with 1 µg RNA and AMV reverse transcriptase (Roboklon) using oligo(dT) primers according to manufacturer’s instructions. qPCR was performed in a volume of 10 µl with the iTaq SYBR GREEN supermix (Biorad) using 45 cycles of 15 sec at 95°C and 30 sec at 60°C in a CFX96 cycler (Biorad). Data were normalized to the transcript encoding PP2A subunit A3 (At1g13320) and expressed as the mean of two biological replicates ± SEM. Primers are listed in Additional file 1.

## Results

### Experimental strategy and evaluation of different probes for specific RNP capture

To enable identification of proteins that interact with a defined mRNA *in vivo*, we adapted the procedure for specific ribonucleoprotein capture developed by Rogell *et al*. for ribosomal RNAs for the use in plants, employing the *AtGRP7* (AT2G21660) mRNA as proof-of-concept [36].

To find the optimal antisense probe, we designed multiple oligonucleotides targeting different regions of the *AtGRP7* transcript (5’UTR, 3’UTR, and exon 2) according to established LNA design guidelines using the QIAGEN LNA Oligo Optimizer tool. Probes were 18-20 nt long and had a melting temperature of ~70-80°C (Fig. 1A). The sequences of the probes were designed to have little self-complementarity, no strong secondary structures, and no complementarity to other transcripts in Arabidopsis (especially to the paralogous *AtGRP8*). Additionally, probes were designed to target regions of the *AtGRP7* transcript that do not contain strong secondary structures, as determined by the Vienna RNAfold web server. At the 3’ end, probes contained a flexible C6 linker and a primary amine, which allows covalent coupling to carboxylated magnetic beads (Fig. 1B). Covalent coupling enables the use of high concentrations of salts and detergents for stringent purification of the RNPs, and the recycling of capture probes, which saves costs. Using the protocol by Rogell *et al*. as a starting point, we systematically optimized the conditions for cell lysis, probe hybridization, removal of unspecific binders and elution for plants.

**Fig. 1:**
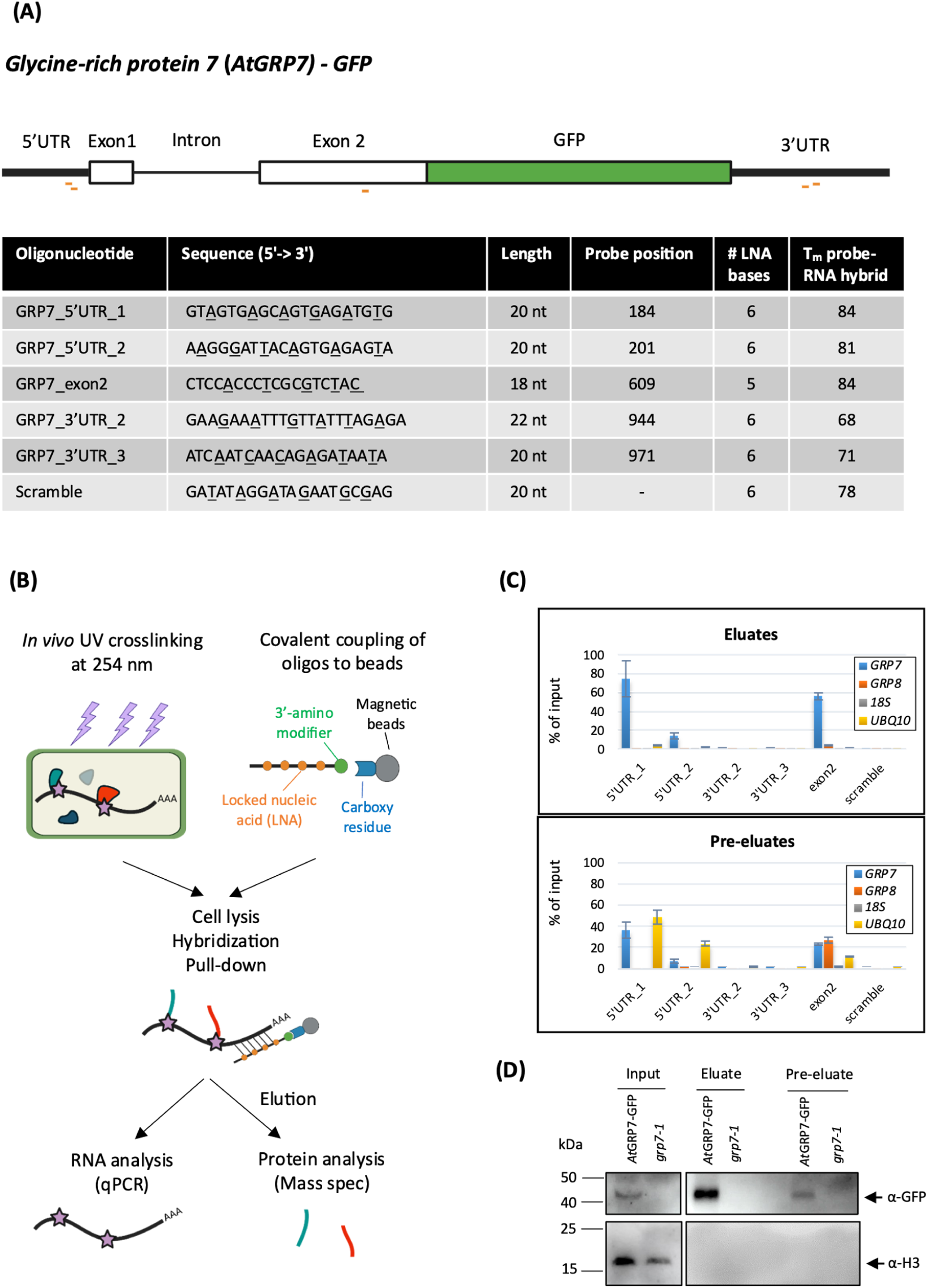
Principle of the *AtGRP7* RNA interactome capture. **(A)** Scheme of the *AtGRP7-GFP* mRNA and details of the antisense LNA/DNA mixmer oligonucleotides tested for capture of the *AtGRP7* transcript. LNA nucleotides are underlined. The probe position is indicated relative to the TAIR cDNA sequence. **(B)** Principle of the specific RNP capture. 14-day-old seedlings are crosslinked with 254 nm UV light to establish covalent bonds between RNA and proteins. The LNA oligonucleotides are coupled to carboxylated magnetic beads via a primary amine attached to a C6 linker at their 3’end. Cell lysates are incubated with the bead coupled oligonucleotides to pull down proteins interacting with the *AtGRP7* RNA. RNA-protein complexes are eluted and either subjected to RNA analysis via RT-qPCR or to protein analysis via mass spectrometry. **(C)** Capture efficiency and specificity with the different LNA/DNA mixmer oligonucleotides. Captures were performed with UV crosslinked *AtGRP7*::*At*GRP7-GFP (*grp7-1)* seedlings and RNA levels of *AtGRP7*, *AtGRP8*, 18S rRNA and *UBIQUITIN 10* were measured in the eluates (top panel) and pre-eluates (bottom panel) by RT-qPCR using the primers listed in Additional file 1. Transcript levels are expressed relative to the transcript level in the input. **(D)** Immunoblot analysis of *AtGRP7*::*At*GRP7-GFP *grp7-1* and *grp7-1* control plant subjected to RNP capture with 5’UTR_1 LNA oligonucleotides. The lysate (input), pre-eluate, and eluate fractions were probed with the α-GFP antibody (top) or the α-Histone H3 antibody (bottom).

As starting material, we used 14-day-old seedlings expressing *At*GRP7-GREEN FLUORESCENT PROTEIN (GFP) under the endogenous promoter in the *grp7-1* background with the rationale to monitor the pulldown of *At*GRP7 known to interact with its own transcript by immunoblotting with the GFP antibody. Seedlings were UV-crosslinked at 2000 mJ/cm^2^ as established for our individual-nucleotide resolution crosslinking and immunoprecipitation (iCLIP) protocol [43, 61]. First, we compared the capture efficiency of the LNA probes directed against different transcript regions (Fig. 1A). To perform the capture, a cell extract was prepared by homogenizing 2 g of plant tissue in 12.5 ml of lysis buffer. The cell extract was then cleared by centrifugation and filtered through a 0.45 µm filter. Probes were coupled to the magnetic beads and heated to 90°C for 3 minutes before hybridization to relax possible secondary structures. Formamide was added to the cell extract to a final concentration of 15% (v/v) to decrease the melting temperature of the probe-RNA duplex and thereby decrease unspecific binding. Hybridization was carried out for 2 h at 4°C under constant rotation. After magnetic separation and removal of the supernatant, beads were washed multiple times followed by a pre-elution step with H_2_O at 40°C for 5 min to remove non-specifically bound transcripts. Finally, RNA-protein complexes were heat-eluted at 90°C for 3 min.

First, we checked for the ability of the oligonucleotides to capture the *AtGRP7* transcript by assessing transcript levels in the input, pre-eluates and eluates using RT-qPCR. The first LNA oligo directed against the 5’UTR (termed 5’UTR_1) clearly performed best, as it captured about 70% of the *AtGRP7* RNA present in the input, while at the same time capturing only small amounts of other transcripts such as *AtGRP8* or the highly abundant *UBQ10* mRNA or 18S rRNA (Fig. 1C). Interestingly, the performance of the second LNA oligo directed against the 5’UTR (termed 5’UTR_2) was considerably worse, even though located in a similar region. The LNA oligo targeting exon2 managed to capture almost 60% of the *AtGRP7* RNA present in the input but was less specific as it also captured more *AtGRP8* transcript. In contrast, the LNA oligos targeting the 3’UTR failed to efficiently capture *AtGRP7* with transcript levels in the eluates not being significantly different to those of the scrambled control. Importantly, the pre-elution step successfully removed unspecific binders (Fig. 1C). Subsequently, we checked whether we pulled down proteins bound to the *AtGRP7* transcript. We performed a capture experiment using the 5’UTR_1 LNA oligo and cell extracts from UV-crosslinked *At*GRP7-GFP plants or *grp7-1* mutants as a control, respectively. Western blot analysis showed that the *At*GRP7 protein, which is known to bind to its own transcript, could be efficiently enriched in the eluate of the *At*GRP7-GFP sample, while being absent in the mutant (Fig. 1D). The abundant DNA-binding protein Histone H3 served as a negative control and was absent in the eluates from both *At*GRP7-GFP and *grp7-1* samples. Together, this demonstrated that the 5’UTR_1 LNA probe is able to efficiently capture *AtGRP7* RNA and associated binding proteins.

### Improved capture efficiency by implementation of tandem capture, optimized buffer conditions and removal of genomic DNA

When transcript levels in the eluate were compared relative to the input, the 5’UTR_1 LNA probe efficiently captured *AtGRP7* RNA (Fig. 1C); however, when comparing levels of different transcripts in the eluate, it becomes apparent that the most abundant RNA species present is 18S rRNA (Fig. 2A). To tackle this issue, we performed a tandem-capture approach similar to the study by [62], where we first used oligo(dT) beads to enrich for all polyadenylated RNAs, followed by specific capture of *AtGRP7* with the 5’UTR_1 LNA probe. Although this could remove some of the 18S rRNA, it still remained the most abundant species in the eluate (Fig. 2B). Interestingly, when we performed the tandem capture the other way around (specific capture followed by oligo(dT)), we could successfully remove the vast majority of ribosomal RNA and other unspecific transcripts, specifically enriching for *AtGRP7* (Fig. 2C).

**Fig. 2:**
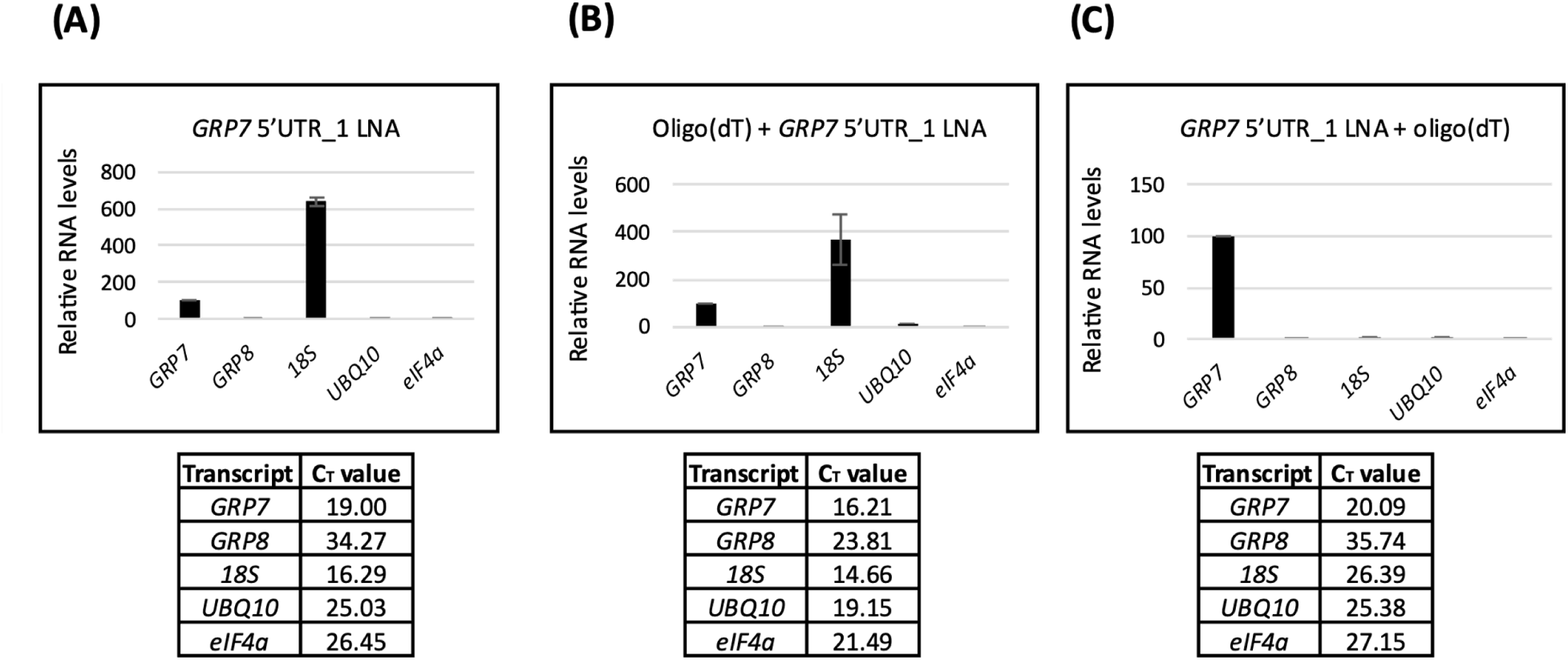
Optimization of the capture efficiency for the LNA 5’UTR_1 probe by tandem capture with oligo(dT). Relative RNA levels in the eluates after a single round of RNP capture with LNA 5’UTR_1 (**A**), tandem capture with oligo(dT) followed by the LNA 5’UTR_1 probe (**B**), and tandem capture with the LNA 5’UTR_1 probe followed by oligo(dT) capture (**C**). The *AtGRP7* level is set to 100%.

However, we noticed some degree of RNA degradation in the lysate, which decreased the amount of transcript present to be captured (Additional file 2). Therefore, we aimed to further optimize the protocol by comparing the lysis and wash buffers [36] that we use here to the ones used in ChIRP-MS [30]. For this comparison, we performed a pulldown employing the ChIRP buffers with small adjustments (Additional file 2). The ChIRP lysis buffer contained NaCl and SDS instead of LiCl and LiDS, and 1 mM EDTA instead of MgCl_2_. Washes were carried out with 2x SSC and 1x SSC buffer, while the wash buffers in Rogell *et al.* (2017) had the same composition as the lysis buffer but with decreasing concentrations of salt and detergents (Additional file 2). Both the RNA integrity in the cell lysate and the capture efficiency as assessed by RT-qPCR could be significantly improved with the ChIRP buffers (Additional file 2). Hence, these buffers were used for all subsequent experiments.

Additionally, the cell lysate was passed once through a 27G needle before probe hybridization to shear genomic DNA (gDNA) and avoid its binding to the probes. RT-PCR of *AtGRP7* using primers that span the intron and can detect both gDNA and mRNA (Additional file 3) showed that only a small amount of gDNA was present in the eluate after capture, even without DNA shearing (Additional file 3). However, inclusion of this step eliminated the last traces of gDNA to a non-detectable level. Moreover, shearing gDNA led to higher levels of captured *AtGRP7* mRNA, while at the same time not increasing the amount of unspecifically bound transcripts (Additional file 3).

### Medium-scale tandem RNA interactome capture only identifies few interacting proteins

After having improved the protocol, we next performed a tandem capture (specific capture with 5’UTR_1 LNA probe followed by oligo(dT)) on a medium scale using 32 g of ground plant material from UV-crosslinked *At*GRP7-GFP plants or the *grp7-1* mutant as a control. Denaturing RNA gel electrophoresis (Fig. 3A) and silver staining (Fig. 3B) showed that RNA and proteins were intact before and after probe hybridization, indicating that there was no degradation during the capture. RT-qPCR analysis demonstrated that *AtGRP7* RNA was specifically enriched over other RNAs in the eluate from the *AtGRP7*-GFP sample and was absent in the *grp7-1* eluate, which contained mostly ribosomal RNA (Fig. 3C). Despite these quality controls looking promising, the MS analysis only returned 18 proteins in total that were quantified (Additional file 4). Of these, only *At*GPR7 had a log_2_ fold-change above 2 compared to the control (Fig. 3D). Gene ontology (GO) term analysis showed that the identified proteins were enriched in RNA-related terms (Fig. 3E), suggesting that the capture protocol *per se* is working, but needed further optimization and up scaling.

**Fig. 3:**
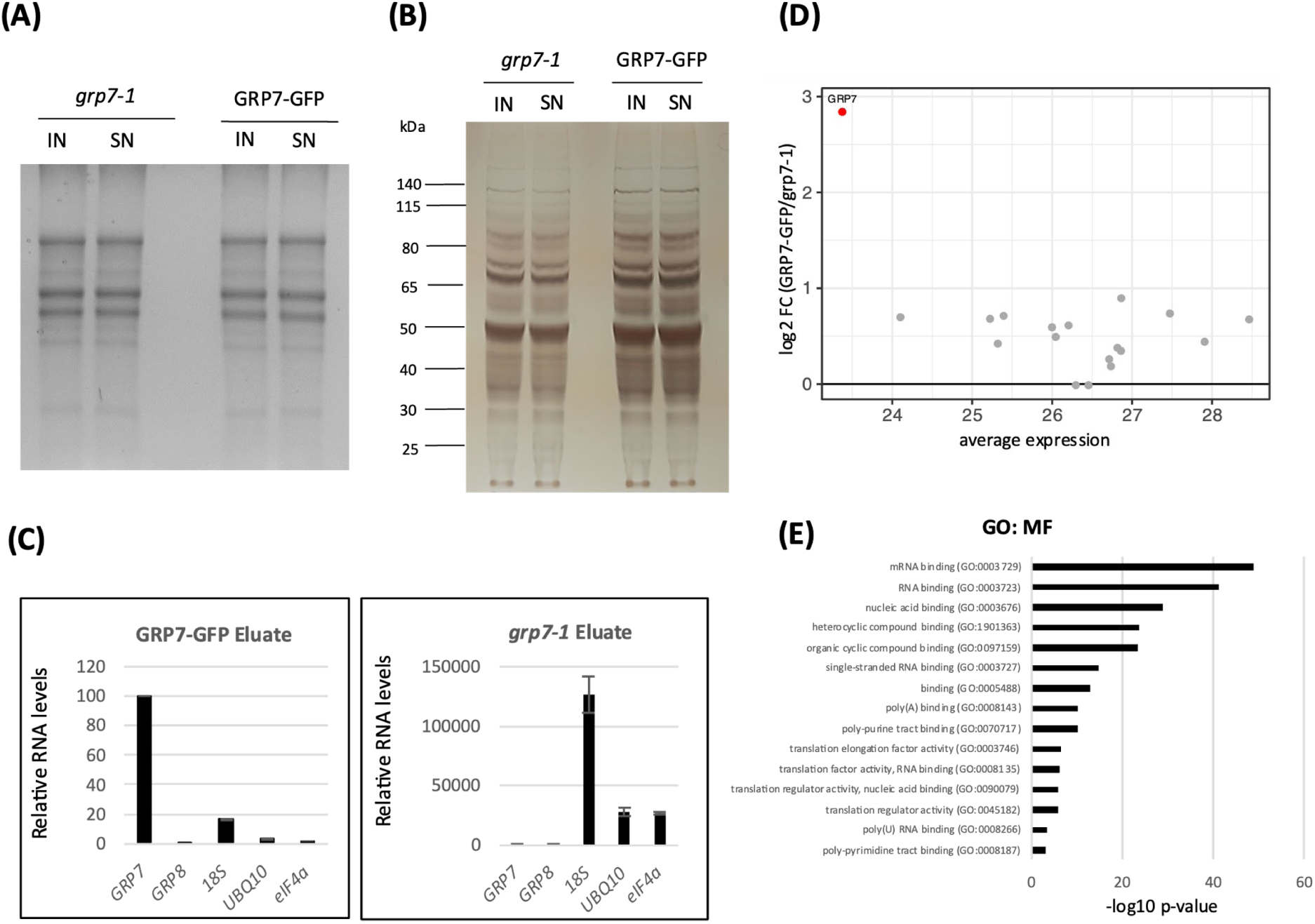
Mass spectrometry analysis of proteins bound to the *AtGRP7* transcript after medium-scale tandem capture. **(A)** Agarose-formaldehyde gel electrophoresis of total RNA in the lysate (input) and the supernatant after probe hybridization (SN) in *AtGRP7*-GFP *grp7-1* plants and *grp7-1* control plants. **(B)** Silver staining of total protein in the lysate (input) and the supernatant after probe hybridization (SN) in the *AtGRP7*-GFP *grp7-1* plants and *grp7-1* control plants. The positions of the molecular weight markers are indicated. **(C)** Relative RNA levels of *AtGRP7*, *AtGRP8*, 18S rRNA, *UBIQUITIN10*, and *eIF4α* RNA in the eluates of *AtGRP7*-GFP *grp7-1* plants (left) and *grp7-1* control plants (right). **(D)** MA plot of identified proteins displaying the relation between log_2_ fold change and the average expression (as log_2_ TMT signal). The *At*GRP7 protein was significantly enriched (red dot). **(E)** GO term analysis of the molecular function of the proteins identified.

### Further optimization by two consecutive captures and replacement of oligo(dT) with LNA2.T probes

To increase the number of RNA-protein complexes in the eluate, we decided to perform two consecutive rounds of capture from the same cell extract with the 5’UTR_1 LNA oligo by using the supernatant after the first hybridization as input for the second round of hybridization with a fresh aliquot of bead-coupled ologonucleotides. Analysis of RNA and proteins after round 1 and round 2 demonstrated that the prolonged time of the protocol did not lead to degradation (Additional file 5). Furthermore, *AtGRP7* RNA could also be efficiently enriched after the second round of capture (Additional file 5).

We also introduced a pre-clearing step with empty beads to reduce unspecific binding. Additionally, we substituted the oligo(dT) beads with 20-mers bearing an LNA-thymine at every other position (LNA2.T), which has been shown to be superior to standard oligo d(T) oligonucleotides for poly(A) RIC [52].

### Identification of *AtGRP7* interacting proteins with optimized large-scale RNA interactome capture

Scaling up the starting material from 32 g to 100 g of UV-crosslinked, ground *At*GPR7-GFP and *grp7-1* tissue, respectively, we performed another tandem capture with two rounds of capture with 5’UTR_1 LNA oligo followed by two rounds of LNA2.T capture, and analyzed the samples by MS. Compared to the previous medium-scale experiment with 32 g of tissue, the total number of quantified proteins was increased to 63, of which 33 had a positive fold change. *At*GRP7 again showed the highest log_2_ fold-change (Fig. 4A). Although the tandem capture has the advantage of specifically enriching for *AtGRP7* RNA in the eluate while removing most of the contaminating transcripts (Additional file 6), much of the *AtGRP7* transcript is lost during this stringent, multi-step procedure with only less than 20% of the input being present in the eluate (Fig. 4B).

**Fig. 4:**
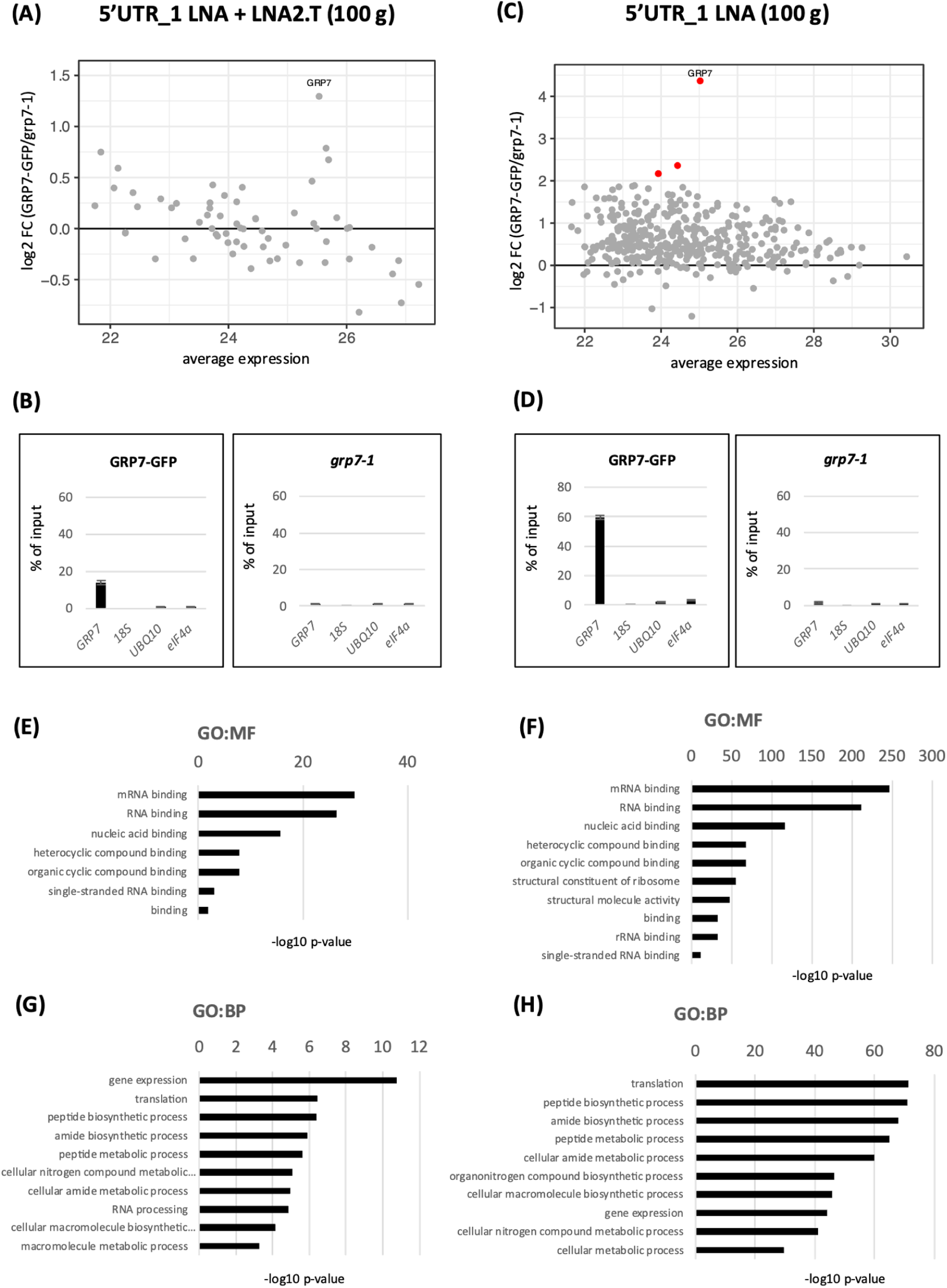
Mass spectrometry of proteins binding to the *AtGRP7* transcript after large-scale capture. MA plot of identified proteins (**A, C**), relative RNA levels of *AtGRP7*, 18S rRNA, *UBIQUITIN 10*, and *eIF4α* in the eluates of *AtGRP7*-GFP *grp7-1* plants (left) and *grp7-1* control plants (right) (**B, D**) and enriched GO terms of identified proteins **(E** and **G, F** and **G**) after large-scale tandem capture with two consecutive rounds of hybridization with the 5’UTR_1 LNA oligo followed by LNA2.T capture, or after large-scale capture with two consecutive rounds of hybridization with the 5’UTR_1 LNA oligo only. Proteins with a log_2_ fold change ≥2) are indicated by the red dots.

Since this is limiting the number of captured proteins, we repeated the large-scale experiment_with 100 g of tissue but did not include the LNA2.T capture. This increased the level of *AtGRP7* transcript in the eluate to about 60% of the input (Fig. 4D), but with the downside of also increasing the amount of nonspecifically bound transcripts, most notably ribosomal RNA (Additional file 6). However, since these unspecific transcripts are also present in the *grp7-1* control (Additional file 6), we reasoned that associated proteins would not be significantly enriched in the MS. Omitting the LNA2.T capture increased the number of proteins to 386 with 356 having a positive fold change and three, including *At*GRP7, having a log_2_ fold-change of at least 2 (Fig. 4C). To validate our MS data, we analyzed the identified proteins from both large-scale (single and tandem) captures by GO term analysis, where we included all proteins with a positive fold change. Although the ones without significant enrichment do not pass the statistical criteria and therefore cannot be clearly distinguished from unspecific binders, they likely contain interesting candidates for further analysis. Indeed, molecular function (MF) GO terms for both large scale captures were linked to RNA-related terms, most notably (m)RNA binding and nucleic acid binding (Fig. 4E-F). Similarly, the top 10 enriched biological process (BP) GO terms were connected to RNA (Fig. 4G-H).

When comparing the proteins with a positive fold change from the large-scale tandem and single RNP captures, we found that almost all proteins (30 out of 33) from the tandem capture were also present in the single capture (Fig. 5A). In this set of common 30 proteins (Additional file 7) almost all of them contained known RNA-binding domains (RBDs), with the RRM domain being the most abundant one (Fig. 5B). Moreover, we performed a STRING network analysis and found that these proteins contain many interaction nodes (Fig. 5C), suggesting that they may be functionally linked in support of authentic interactions. Taken together, these results show that even though we find only few proteins enriched upon pulldown in *At*GRP7-GFP plants relative to the *grp7-1* control, our *in vivo* approaches using LNA oligonucleotides were nevertheless able to strongly enrich for proteins know to be involved in RNA binding and potentially function as novel regulators of *AtGRP7*.

**Fig. 5:**
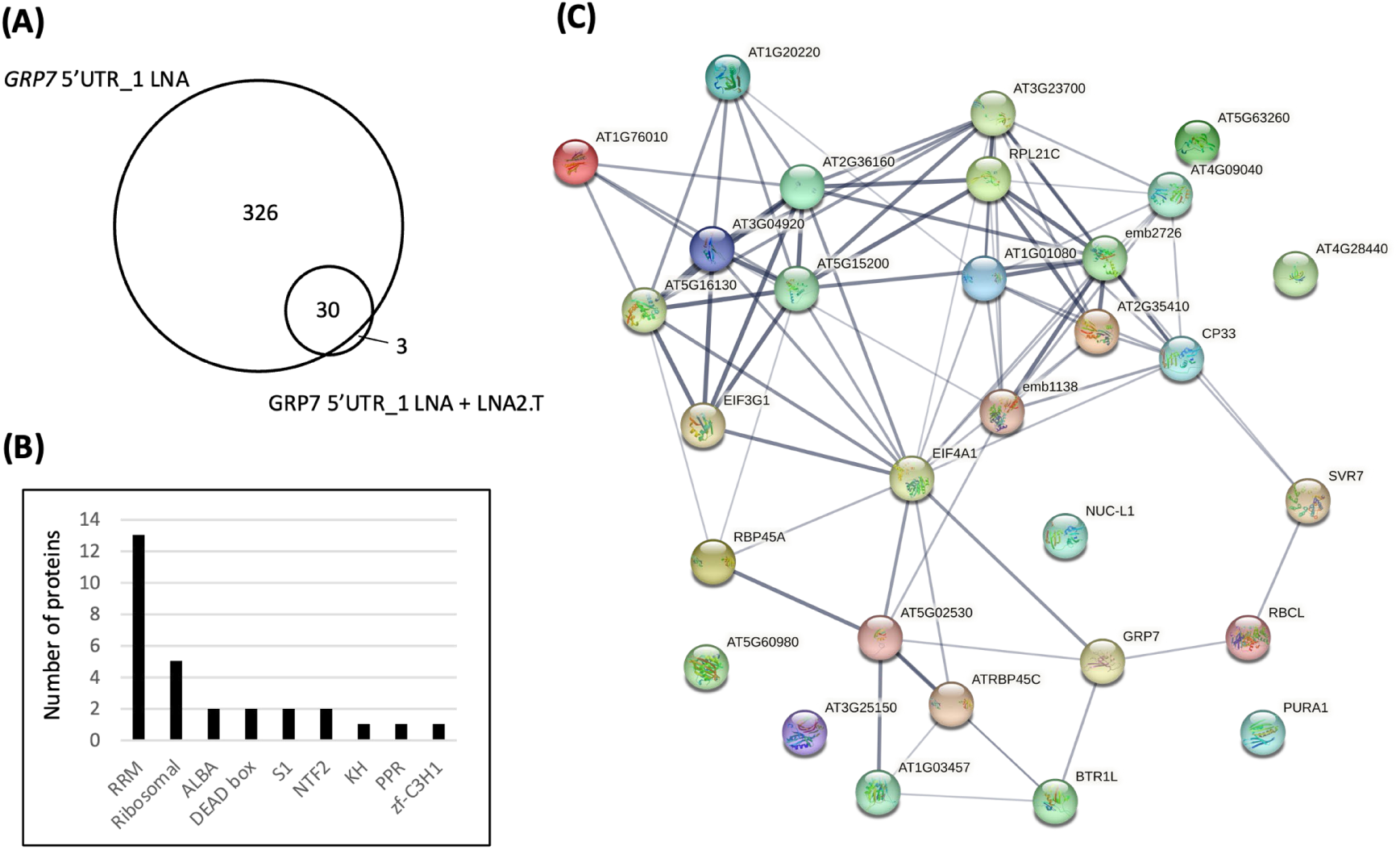
Properties of proteins with a positive fold change. **(A)** Venn diagram showing the overlap between the proteins identified in the large-scale capture with two consecutive rounds of hybridization with the 5’UTR_1 LNA oligo and the large-scale tandem capture with hybridization with 5’UTR_1 LNA oligo followed by LNA2.T capture. **(B)** Number of proteins annotated with classical RNA-binding domains among the 30 common proteins. (**C)** STRING network analysis of the 30 common proteins.

### Identification of proteins interacting with *AtGRP7* UTRs by *in vitro* RNA interactome capture

The proteins recovered by specific capture of *At*GRP7 *in vivo* were compared to a set of proteins identified by *in vitro* pulldowns of the *AtGRP7* 5’UTR and 3’UTR. These regions were chosen because in iCLIP experiments we obtained extensive crosslinking of the *At*GRP7 protein to these regions in iCLIP [43].

As a bait, we used *in vitro* transcribed, biotinylated RNA of the *AtGRP7* 5’UTR and 3’UTR, respectively, coupled to magnetic Dynabeads MyOne Streptavidin beads. The 3’UTR RNA bait contained the annotated 3’UTR sequence, and the 5’UTR RNA bait contained the 5’UTR sequence including the transcription start site [44] and 15 nucleotides of the first exon (Additional file 8). These bait RNAs were incubated with nucleoplasmic extracts prepared from 5 g of ground 14-day old Col-0 wild-type plants to avoid contamination with the highly abundant DNA-binding proteins coming from the chromatin. Unspecific background was minimized by washing the immobilized RNA–protein complexes three times with buffer containing 100 mM sodium chloride and 0.1% Triton X-100 as a non-ionic detergent. RNA-protein complexes were then recovered by heat-elution in LDS sample buffer. A pulldown with empty beads served as negative control.

We first performed a test pulldown to check RNA integrity after different steps of the protocol. As demonstrated by urea PAGE, the 5’UTR and 3’UTR bait RNAs were efficiently coupled to the beads, as they were depleted in the supernatant (SN) after coupling (Fig. 6A, 6B). Importantly, both bait RNAs were intact before and after the pulldown (Fig. 6A, 6B). Silver staining showed that our *in vitro* capture approach could enrich for specific proteins as compared to the empty beads control (Fig. 6C, 6D). We then performed four biological replicates for each capture and analyzed the proteins by MS. Capture with the 5’UTR and 3’UTR RNA baits identified 213 and 356 proteins, respectively, that were significantly enriched (log_2_FC ≥ 1; p-value ≤ 0.05) over empty beads (Fig. 6E, 6F). For both pulldowns, the GO terms of these proteins were strongly enriched in RNA-related terms, validating our *in vitro* approach (Additional file 9).

**Fig. 6:**
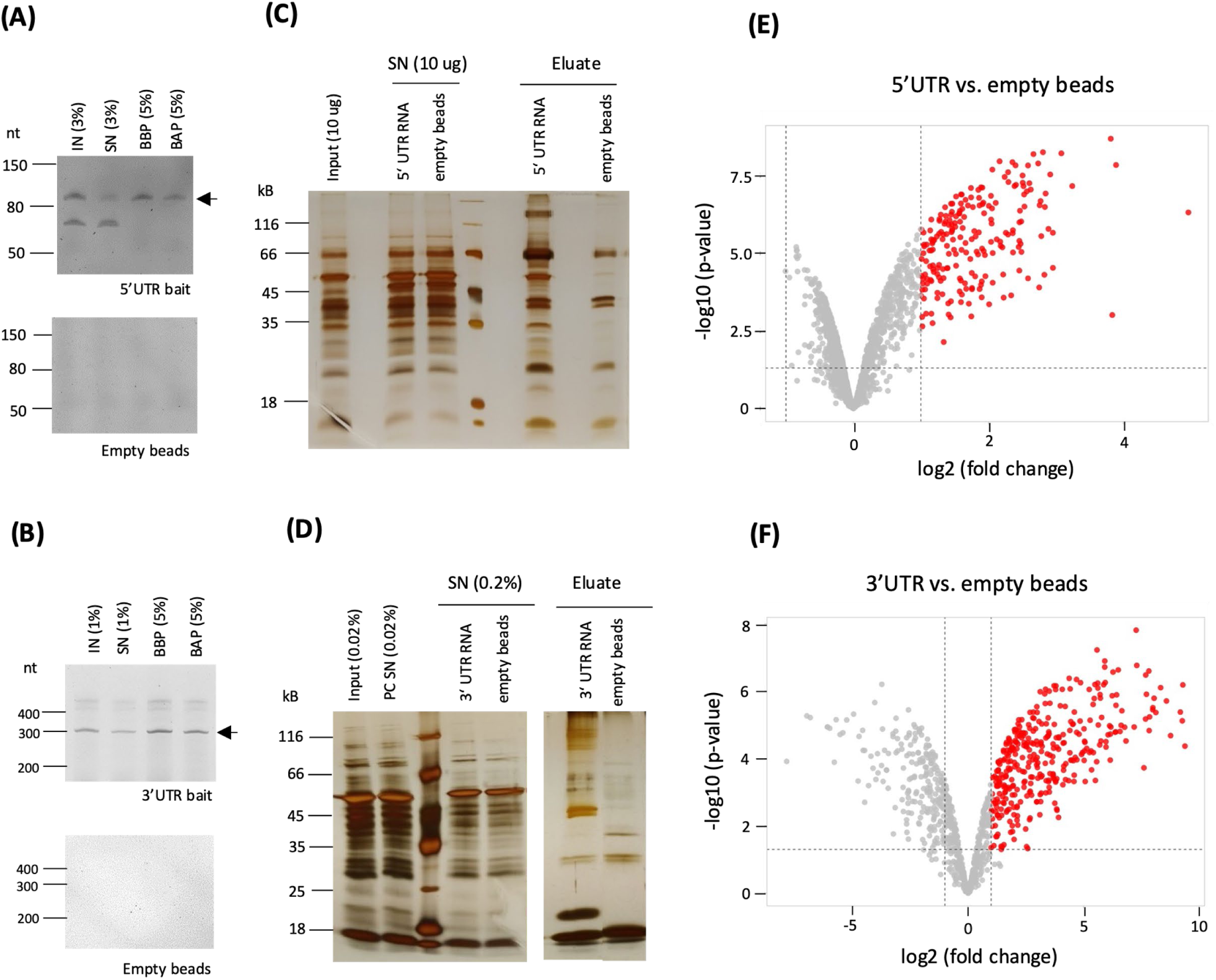
Identification of *AtGRP7* binding proteins by *in vitro* pulldowns. **(A, B)** Coupling efficiency of biotinylated 5’UTR bait (**A**) and 3’UTR bait (**B**) to magnetic streptavidin beads. Aliquots of the RNA isolated from the input (IN), supernatant after coupling (SN) and eluates from the beads before (BBP) and after pulldown (BAP) were analyzed on 12.5% urea PAGE gels. Empty beads were used as controls. The arrows indicate the *in vitro* transcripts of the sizes expected for the 5’UTR (**A**) or 3’UTR (**B**), respectively. The additional bands likely represent additional conformations. **(C, D)** Silver staining of proteins recovered after *in vitro* captures with 5’UTR bait (**C**) and 3’UTR bait (**D**). Aliquots of the input, supernatant and eluates of the respective coupled and empty beads were separated on a 12% SDS-Page and subjected to silver staining. **(E, F)** Volcano plots of proteins identified by *in vitro* capture with the 5’UTR bait (**E**) or 3’UTR bait (**F**). Significantly enriched proteins (log_2_ fold change ≥1, p-value <0.05) are indicated by the red dots.

Next, we compared the significantly enriched proteins from the *in vitro* pulldowns to the proteins with a positive fold change from the *in vivo* large-scale capture with the *AtGRP7* 5’UTR_1 LNA oligo (without subsequent LNA2.T capture). 59 and 71 proteins overlapped between the *in vivo AtGRP7* LNA large-scale capture and the *in vitro* 5’UTR and 3’UTR capture, respectively, with 44 proteins being common to all three captures (Additional file 10).

### Identification of novel regulators of *AtGRP7*

To identify potential novel regulators of the *AtGRP7* transcript, we focused on proteins with classical RBDs, which were found in more than one capture. The most abundant of these were proteins with an RRM, of which we identified 40 in total that were present in at least two of the capture experiments (Fig. 7). Moreover, we found several nuclear transport factors, RNA helicases and KH domain containing proteins (Fig. 7) in addition to several other protein groups related to RNA processing, such as translation factors, poly(A) binding proteins and small RNA related proteins (Additional file 10). But most strikingly, we identified five out of the six ALBA (Acetylation Lowers Binding Affinity) proteins (Fig. 7). ALBA proteins are highly conserved and found in all kingdoms of life [63]. Despite their strong conservation, they appear to have diverse functions. In archaea, ALBA proteins are mostly involved in genome packaging and organization [64], while in protozoa, they regulate RNA stability and translational [65]. In yeast and humans, ALBA proteins are involved in tRNA processing [66]. Arabidopsis encodes six ALBA proteins. ALBA1, ALBA2 and ALBA3 are short proteins, which almost solely consist of the ALBA domain, while ALBA4, ALBA5 and ALBA6 contain additional arginine-glycine (RGG) repeats at their C-termini, which mediate nucleic-acid binding and protein-protein interactions [67]. ALBA1 and ALBA2 were reported to be localized in both the nucleus and the cytoplasm. In the nucleus, they bind to genic R-loops to maintain genome stability [68]. ALBA4-6 are functionally redundant, as only *alba456* triple mutants, but not single or double mutants, show pleiotropic developmental defects [49, 69]. Under heat stress, ALBA4-6 were shown to bind to selected transcripts and recruit them into phase separated stress granules and processing bodies for protection [69].

**Fig. 7:**
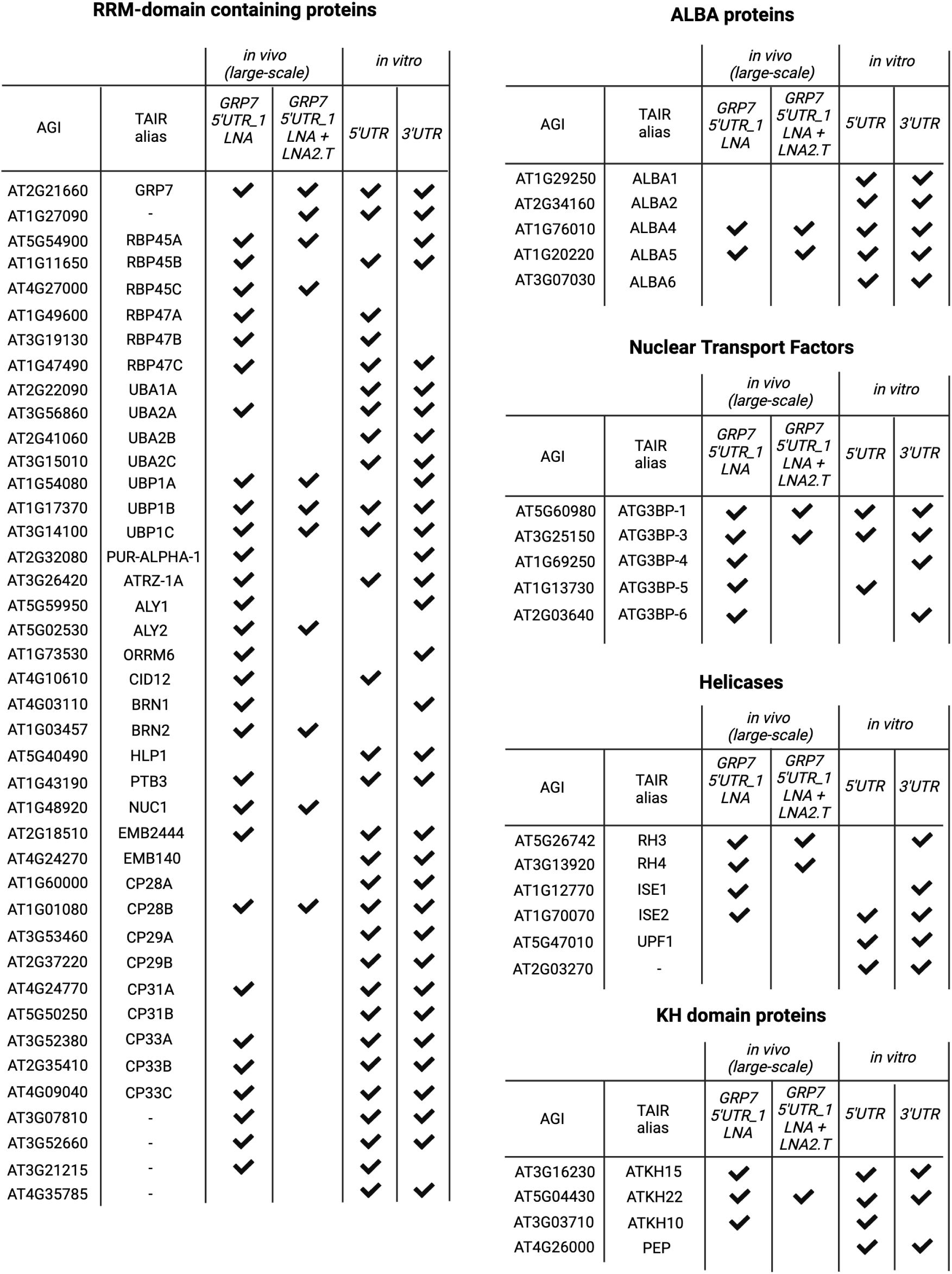
Overview of prominent protein groups identified by *in vivo* and *in vitro* pulldowns of *AtGRP7* interactors.

Recently, the direct *in vivo* target transcripts of ALBA4 were determined on a global scale via individual nucleotide resolution crosslinking and immunoprecipitation (iCLIP) using ALBA4-GFP lines in an *alba456* background that will be reported elsewhere (Reichel et al., manuscript in preparation). Plants expressing GFP under the 35S promoter were used as control. Analyzing this dataset regarding the interaction between ALBA4 and *AtGRP7*, we found many binding sites in both the *AtGRP7* 5’- and especially the 3’UTR (Fig. 8), independently validating our MS data.

**Fig. 8:**
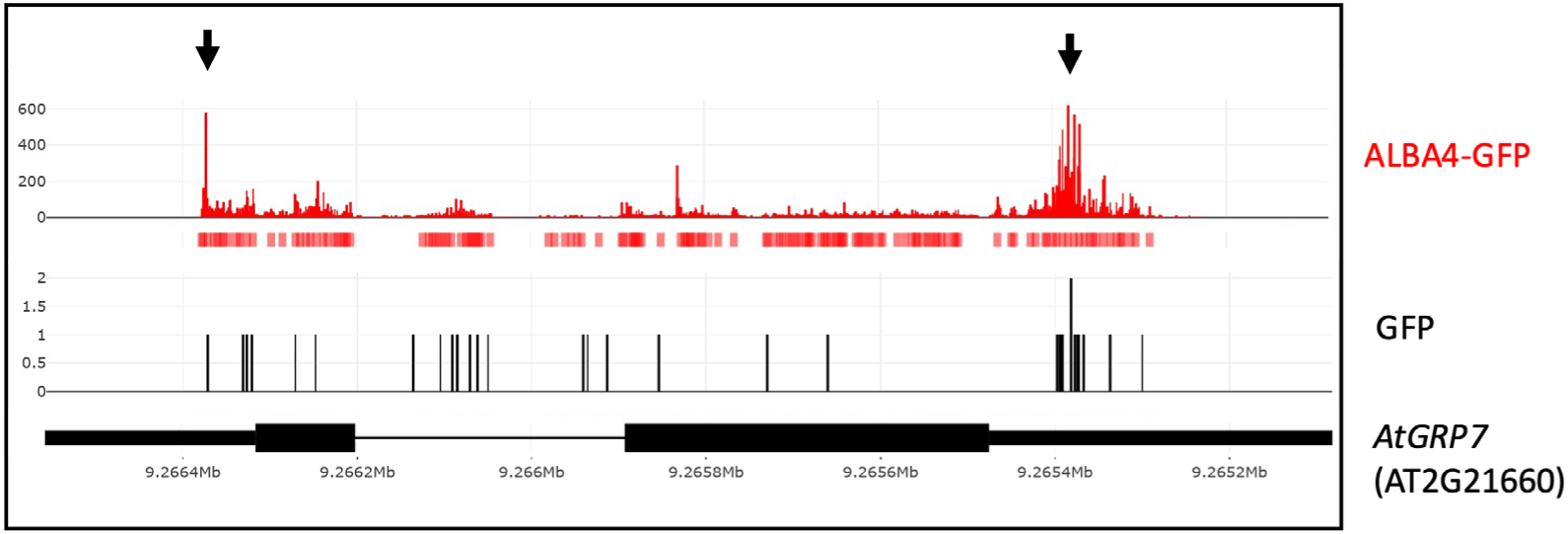
*In vivo* binding of ALBA4 to the *AtGRP7* transcript. Binding sites of *ALBA4*::ALBA4-GFP in *alba456* to *AtGRP7* determined by iCLIP (Reichel et al., manuscript in preparation). iCLIP peaks of ALBA4 are shown in red and peaks of a GFP control sample are shown in black. The red boxes below the ALBA4-GFP iCLIP reads denote called peaks. Prominent ALBA4 peaks are highlighted by arrows. The *AtGRP7* gene model is shown at the bottom. Black boxes: exons; narrow black boxes: untranslated regions; line: intron.

To test for a functional relevance of ALBA4 binding to the *AtGRP7* transcript, we performed an extensive time course analysis in the *alba456* mutant compared to Col-0 wild type plants. Seedlings were grown in entraining long day (LD) conditions, shifted to continuous light (LL) and harvested at 2 h intervals, starting at LL28. *AtGRP7* oscillates in wild type plants with a peak at the end of the daily light period, and the oscillations persist in LL. In LD conditions, there is no difference in oscillation between Col-0 and *alba456* (Fig. 9A). In LL conditions however, *AtGRP7* levels reach the maximum about 2 h earlier in *alba456* than in Col-0, which becomes most apparent on day three in LL (Fig. 9B). Moreover, *AtGRP7* levels rise and decline faster in *alba456* than in Col-0. Even though the changes are subtle, this indicates that ALBA proteins influence the oscillation of the *AtGRP7* transcript in extended light conditions.

**Fig. 9:**
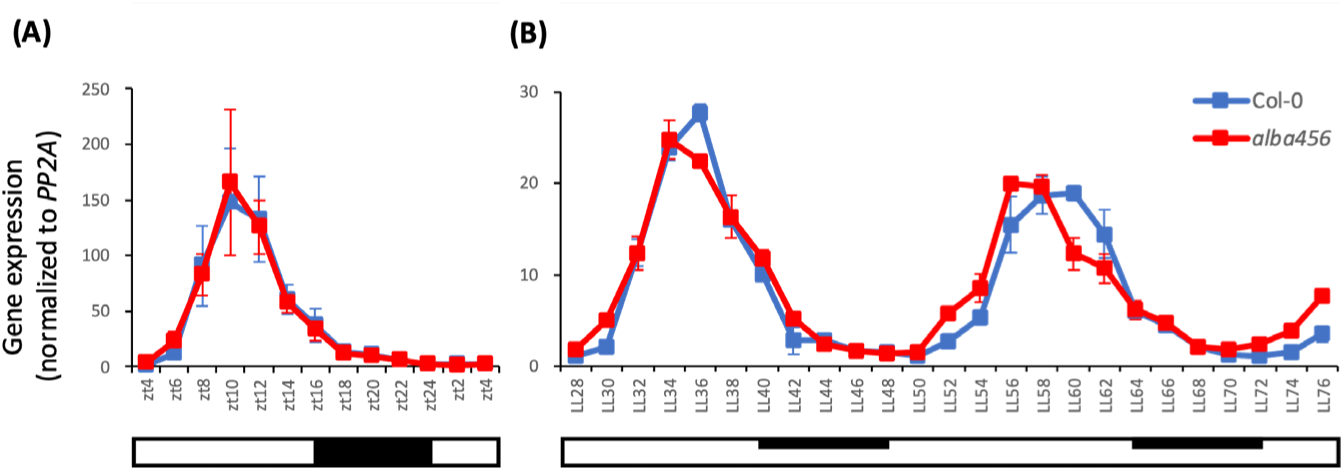
Impact of altered ALBA protein levels on *AtGRP7* transcript oscillations. Col-0 wild type plants and the *alba456* triple mutant were grown in long day conditions (16h light/8h dark) for 12 days before transfer to constant light (LL). Seedlings were harvested at 2-h intervals throughout the light-dark cycle (**A**) and from 28 h to 76 h in LL (**B**). *AtGRP7* transcript levels were analyzed by RT-qPCR and normalized to *PP2A*. Error bars represent the standard deviation of two biological replicates. Open bar: constant light; dark bar: (subjective) night.

## Discussion

Identification of the protein binding partners of a single mRNA is vital for understanding how its mRNA is regulated. So far, most studies have focused on abundant non-coding RNAs or viral RNAs and were performed in mammalian cell cultures [28, 30, 32, 36, 70, 71]. More recently, identification of proteins that bind to a germ-line specific transcript in *Caenorhabditis elegans* has been reported [72]. Here, we have for the first time determined the protein-binding repertoire of a single plant mRNA *in vivo*, using the clock regulated *AtGRP7* transcript as proof-of-concept. Towards this end, we combined and optimized elements of different RIC protocols.

Several aspects have proven critical for the success of the approach: firstly, the design of antisense oligonucleotides is key for efficient recovery of target mRNAs. We recommend testing multiple probes that bind to different regions in the transcript to optimize specificity and efficacy. In parallel to optimizing the pulldown for LNA oligonucleotides, we also compared a combination of ten antisense DNA oligonucleotides, as used in ChIRP for non-coding RNAs, that were tiled across the entire *AtGRP7* transcript (sequences listed in Additional file 11). Notably, LNA antisense oligonucleotides outperformed tiled DNA oligonucleotides that were not able to enrich for the *AtGRP7* transcript in our conditions (Additional file 11). This could be due to coverage by translating ribosomes.

Secondly, an optimal UV crosslinking dosage is important to maximize recovery of binding proteins. For 14-day-old seedlings, we chose a dosage of 2000 mJ/cm^2^, which has proven optimal in protein centric CLIP methods [18]. However, if another type of tissue is used, we recommend testing different crosslinking energies. At the same time, it is important to assess RNA integrity in the crosslinked sample as over-exposure can lead to RNA degradation and hence decreased mRNA recovery. The inefficient UV crosslinking, which is even more pronounced in plants due to the presence of UV-absorbing pigments like chlorophyll, combined with a limited capture efficacy of about 60% of the input makes the retrieval of specific protein interactors challenging.

Thirdly, stringent controls are necessary to discriminate specific binders from background, when working with whole plants which increases tissue complexity and unspecific binders. We opted for the mutant *grp7-1* lacking the transcript of interest, which controls for background inherent to the experimental setup. Alternatively, one could use a non-crosslinked sample or perform a pull down with scrambled oligonucleotides.

Even though we could improve the protocol for use in plants by optimizing lysis and washing buffers, shearing genomic DNA, performing two rounds of capture and considerable up-scaling to 100 g of starting material, we still detected only a small number of proteins that were significantly enriched.

Recently, a tandem RNA isolation procedure (TRIP) involving mRNA isolation followed by capture with 3’-biotinylated 2’-O-methylated RNA AOs was used to capture mRNA-protein complexes from yeast, *C. elegans* and HEK293 cells [62]. When we applied a tandem capture approach using the specific GRP7 5’UTR_1 LNA oligo first, followed by capture with LNA2.T oligonucleotides, it resulted in specific recovery of the *AtGRP7* mRNA and removal of background binders, although the number of identified proteins was small. Omitting the LNA2.T capture increased the number of identified proteins considerably, but at the cost of also capturing more unspecific binders. Nevertheless, MF GO terms for both large scale captures were linked to RNA-related terms, most notably (m)RNA binding and nucleic acid binding. Similarly, the top 10 enriched BP GO terms were connected to RNA, highlighting the value of the optimized protocol to recover interactors with a potential role in RNA regulation.

The 356 proteins identified in the large captures with two rounds of 5’UTR_1 LNA oligo were benchmarked against proteins recovered by an *in vitro* pulldown of nucleoplasmic proteins with either the *AtGRP7* 3’UTR or 5’UTR as a bait (Additional file 9). 59 of the *in vivo* identified proteins overlapped with proteins binding to the 5’UTR *in vitro*, and 71 overlapped with proteins binding to the 3’UTR *in vitro*, providing evidence for binding to the *AtGRP7* transcript. Amongst the common proteins between the *in vivo* and *in vitro* approaches were the ALBA protein family. The *AtGRP7* transcript was independently verified as target of ALBA4 by a recent iCLIP experiment, which determined strong ALBA4 binding sites in the *AtGRP7* 5’UTR and especially in the 3’UTR (Reichel et al., manuscript in preparation). By performing an extensive time course experiment, we showed that the *AtGRP7* transcript reaches its expression peak earlier in the *alba456* mutant compared to Col-0 in extended light conditions, suggesting that ALBA proteins alter the oscillation of *AtGRP7*. Although the observed effect is rather mild under standard growth conditions, it is possible that there is a more pronounced effect under stress, since both *At*GRP7 and ALBA proteins are involved in stress response [69, 73, 74].

## Conclusion

We adapted specific RNA interactome capture for use in plants, using *AtGRP7* as a showcase, and identified the ALBA protein family as new regulators. Since *AtGRP7* is one of the most abundant mRNAs at its expression peak, the protocol will still need optimization and up-scaling to be applicable for other less abundant mRNAs and to cover the whole spectrum of interacting proteins. To increase the yield of recovered proteins, this protocol could be used with a more potent cross-linking reagent, like formaldehyde, instead of UV light. However, this would have the disadvantage of recovering both direct and indirect interactors. Continuous improvements of mass spectrometry sensitivity will also help to increase the range of identified proteins.

## Abbreviations

ALBA: ACETYLATION LOWERS BINDING AFFINITY
AO: antisense oligonucleotide
ASCO: ALTERNATIVE SPLICING COMPETITOR
AtGRP7: *Arabidopsis thaliana* GLYCINE RICH RNA-BINDING PROTEIN 7
BAP: beads after pulldown
BBP: beads before pulldown
CHART: Capture Hybridization Analysis of RNA Targets
EDC-HCl: *N*-(3-dimethylaminopropyl)-*N′*-ethylcarbodiimide hydrochloride
eRIC: enhanced RNA interactome capture
gDNA: genomic DNA
GFP: Green fluorescent protein
GO: Gene ontology
h: hour
iCLIP: individual-nucleotide resolution crosslinking and immunoprecipitation
IN: input
LD: light-dark
LL: continuous light
LNA: locked nucleic acid
lncRNA: long noncoding RNA
MALAT1: metastasis-associated lung adenocarcinoma transcript 1
MES: 2-(N-morpholino)ethanesulfonic acid
min: minute
MS: Mass spectrometry
NEAT1: nuclear-enriched abundant transcript 1
NSRa: NUCLEAR SPECKLE RNA-BINDING PROTEIN a
PBS: phosphate buffered saline
PCR: Polymerase chain reaction
PHAROH: Pluripotency and Hepatocyte Associated RNA Overexpressed in HCC
PVP40: polyvinylpyrrolidone 40
PWB: protein wash buffer
RBPome: RNA binding proteome
RIC: RNA interactome capture
RNP: Ribonucleoprotein
RT: reverse transcription
SN: supernatant
snRNA: small nuclear RNA
UTR: untranslated region
UV light: ultraviolet light

## Supplementary information

**Additional file 1:** Primers used in this study.

**Additional file 2: Experimental procedure and composition of the buffers used for optimization of the specific RNP capture.**

**(A, B)** Buffer composition and protocol based on Rogell *et al*. (2017) for specific capture of rRNA binding proteins **(A)** and Chu *et al*. (2015) for ChIRP-MS **(B)**. **(C, D)** Analysis of RNA levels in eluates determined by RT-qPCR (left), corresponding C_T_ values (middle), and agarose-formaldehyde gel electrophoresis of total RNA in the input of crosslinked and non-crosslinked samples (right) after tandem capture with the LNA 5’UTR_1 probe followed by oligo(dT) capture using the protocol of Rogell *et al.*, 2017 **(C)** or Chu *et al*., 2015 **(D)**.

**Additional file 3: Removal of residual genomic DNA by shearing.**

**(A)** Schematic diagram of primers spanning the *AtGRP7* intron and amplicons from genomic DNA (gDNA), alternatively spliced (AS) pre-mRNA and mature mRNA. **(B)** Comparison of the level of genomic *At*GRP7 DNA in the cell lysate without passage (−27G needle) or after passage through a 27G needle (+27G needle). RT-PCR was performed with (+RT) or without (−RT) prior reverse transcription. For comparison, amplification from genomic DNA (gDNA) is shown. Arrows indicate the amplicons derived from gDNA, alternatively spliced pre-mRNA and the fully spliced mRNA [42]. **(C)** Relative levels of *AtGRP7*, *AtGRP8*, *UBIQUITIN 10*, and *eIF4α* mRNA (blue) or gDNA contamination (orange) in eluates without passage (−27G needle) or after passage of the lysate through a 27G needle (+27G needle). The *AtGRP7* level is set to 100 %.

**Additional file 4: MS analysis**

Differential abundance analysis of proteins identified by mass spectrometry

**Additional file 5: Protein and RNA analysis upon two rounds of capture with 5’UTR_1 LNA oligo.**

**(A)** Agarose-formaldehyde gel electrophoresis of total RNA in the lysate (input) and the supernatant after the first round (SN1) and second round (SN2) of probe hybridization in the *AtGRP7*-GFP *grp7-1* plants. **(B)** Silver staining of total protein in the lysate (input) and the supernatant after the first round (SN1) and second round (SN2) of probe hybridization in the *AtGRP7*-GFP *grp7-1* plants. The positions of the molecular weight markers are indicated. **(C)** Relative *AtGRP7*, *AtGRP8*, 18 S rRNA, *UBIQUITIN10*, and *eIF4α* RNA levels in the eluates of the first round (top) and the second round (bottom). Right, corresponding CT values.

**Additional file 6: RNA levels in the eluates after the large-scale capture.**

Relative *AtGRP7*, 18S rRNA, *UBIQUITIN 10*, and *eIF4α* RNA levels in the eluates of *AtGRP7*-GFP *grp7-1* plants (left) and *grp7-1* control plants (right) upon large scale tandem capture with two consecutive rounds of hybridization with the 5’UTR_1 LNA oligonucleotide followed by LNA2.T capture **(A)**, or large-scale capture with two consecutive rounds of hybridization with the 5’UTR_1 LNA oligo only **(B)**.

**Additional file 7**: **List of 30 common proteins** from the large-scale capture with two consecutive rounds of hybridization with the 5’UTR_1 LNA oligonucleotide and the large-scale tandem capture with hybridization with 5’UTR_1 LNA oligonucleotide followed by LNA2.T capture.

**Additional file 8**: **Sequences of baits** derived from the *AtGRP7* 5’UTR **(A)** and 3’UTR **(B)** used for *in vitro* pulldowns.

**Additional file 9: GO term analysis.**

Enriched GO terms for molecular function **(A)** and biological process **(B)** for proteins significantly enriched after *in vitro* capture with the 5’UTR bait (left) and 3’UTR bait (right). **(C)** Venn diagram of proteins identified by large scale *in vivo* capture with the 5’UTR_1 LNA oligo and significantly enriched proteins from the *in vitro* pulldowns.

**Additional file 10**: **Overview of additional groups of proteins identified by *in vivo* and *in vitro* pulldowns of *AtGRP7* interactors.**

Additional file 11: Capture with antisense DNA oligonucleotides

**(A)** Sequences of antisense (AS) DNA oligonucleotides directed against *AtGRP7*. **(B)** Relative *AtGRP7*, 18S rRNA, *UBIQUITIN 10*, and *eIF4α* RNA levels in the eluates of *AtGRP7*-GFP *grp7-1* plants upon capture with AS DNA oligonucleotides or with the 5’UTR_1 LNA oligo. **(C)** Agarose-formaldehyde gel electrophoresis of total RNA in the lysate (input) and the supernatant (SN) after hybridization with AS DNA oligonucleotides or the 5’UTR_1 LNA oligo in *AtGRP7*-GFP *grp7-1* plants.

## Acknowledgements

We thank Elisabeth Klemme and Kristina Neudorf for expert technical assistance and Tony Millar and Naiqi Wang (ANU, Canberra) for providing the ALBA4-GFP and 35S-GFP lines. We thank Dr. Zhicheng Zhang for valuable comments on the manuscript.

Figures were created with Biorender.

## Availability of data and materials

The mass spectrometry proteomics data have been deposited to the ProteomeXchange Consortium via the PRIDE [75] partner repository with the dataset identifiers PXD048975 and PXD049303.

## Declarations

### Ethics approval and consent to participate

Not applicable

### Consent for publication

Not applicable

### Authors’ contributions

D.S. and M.Reichel performed project conception and design.

M.Reichel performed *in vivo* pulldowns. O.S. performed *in vitro* pulldowns.

M. Rettel and F.S. performed proteomics analysis of the in vivo pullodwns and 5’UTR in vitro pulldowns. F.B. performed proteomics analysis of the 3’UTR in vitro pulldowns.

T.K. provided advice on pulldowns.

M. Reichel and D.S. wrote the main manuscript text. M.Reichel prepared the figures.

All authors reviewed the manuscript.

